# Depletion of cardiolipin induces major changes in energy metabolism in *Trypanosoma brucei* bloodstream forms

**DOI:** 10.1101/2020.06.15.152785

**Authors:** Mauro Serricchio, Carolina Hierro-Yap, David Schädeli, Hisham Ben Hamidane, Andrew Hemphill, Johannes Graumann, Alena Zíková, Peter Bütikofer

**Affiliations:** Institute of Biochemistry and Molecular Medicine, University of Bern, Bern, Switzerland; Faculty of Science, University of South Bohemia, Ceske Budejovice, Czech Republic; Graduate School for Cellular and Biomedical Sciences, University of Bern, Bern, Switzerland; Weill Cornell Medicine - Qatar, Doha, State of Qatar; Institute of Parasitology, Vetsuisse Faculty, University of Bern, Bern, Switzerland; Institute of Parasitology, Biology Centre, Czech Academy of Sciences, Ceske Budejovice, Czech Republic

**Keywords:** Mitochondria, cardiolipin, electron transport chain, protein complexes, trypanosomes, ATP synthase

## Abstract

Cardiolipin (CL) is a mitochondrial inner membrane glycerophospholipid that associates with mitochondrial proteins to promote their activities and to facilitate protein complex and super-complex formation. Loss of CL leads to destabilized respiratory complexes and mitochondrial dysfunction. The role of CL in an organism lacking a conventional electron transport chain (ETC) has not been elucidated so far. We now report that in *Trypanosoma brucei* bloodstream forms, in which the ETC is truncated and composed of alternative oxidase and glycerol-3-phosphate dehydrogenase, and the mitochondrial membrane potential is generated by the hydrolytic action of the F_o_F_1_-ATP synthase, the inducible depletion of cardiolipin synthase (TbCls) is essential for parasite survival. Loss of TbCls and CL caused a rapid drop in ATP levels and a decline in the mitochondrial membrane potential. Unbiased proteomic analyses revealed a reduction in the levels of many mitochondrial proteins, most notably of F_o_F_1_-ATP synthase subunits and of the alternative oxidase, resulting in a strong decline of glycerol-3-phosphate-stimulated oxygen consumption. Interestingly, the changes in cellular respiration preceded the observed decrease in F_o_F_1_-ATPase stability, suggesting that the truncated ETC is the first pathway responding to the decline in CL. In addition, proteomic and metabolomic analyses revealed that select proteins and pathways involved in glucose and amino acid transport and metabolism are up-regulated during CL depletion, possibly as a stress response to restore cellular ATP levels.

## Introduction

Cardiolipin (CL) is a mitochondrial and bacterial glycerophospholipid consisting of four fatty acyl chains and two phosphate groups. Owing to its structure, CL has biophysical properties that set it apart from other glycerophospholipids (reviewed by (Schlame & Ren, 2009)). CL adopts a hexagonal phase and thus localizes preferentially to sites with high membrane curvature, such as bacterial septa and poles (Mileykovskaya & Dowhan, 2000; Renner & Weibel, 2011), and it self-organizes in negatively curved membranes (Beltran-Heredia et al., 2019) found at cristae junctions (Friedman, Mourier, Yamada, McCaffery, & Nunnari, 2015) and cristae tips (Acehan et al., 2011). CL tightly associates with respiratory complexes (Fiedorczuk et al., 2016; Lange, Nett, Trumpower, & Hunte, 2001; Shinzawa-Itoh et al., 2007; Solmaz & Hunte, 2008) and is required for the assembly and stability of super-complexes (Mileykovskaya & Dowhan, 2014) and the oligomerization of ATP synthase (Acehan, et al., 2011). In addition, several mitochondrial carriers have been shown to interact with CL (Claypool, 2009; Klingenberg, 2009). Finally, CL promotes mitochondrial membrane fusion (Joshi, Thompson, Fei, Huttemann, & Greenberg, 2012), iron-sulfur cluster biogenesis (Patil, Fox, Gohil, Winge, & Greenberg, 2013) and induction of apoptosis (Santucci, Sinibaldi, Cozza, Polticelli, & Fiorucci, 2019).

More recently, depletion of CL has been shown to affect a cell’s energy metabolism. In a *Saccharomyces cerevisiae* mutant lacking CL, due to deletion of the *crd1* gene encoding CL synthase, production of acetyl-CoA was decreased as a result of a defect in the pyruvate dehydrogenase bypass pathway (Li et al., 2019). This, together with the observation that CL is required for the proper function of the iron-sulfur-containing enzymes aconitase and succinate dehydrogenase (Patil, et al., 2013), demonstrated that the tricarboxylic acid (TCA) cycle is impaired in CL-deficient yeast. As a result, in this *crd1*Δ mutant anaplerotic pathways are activated to restore acetyl-CoA and TCA cycle intermediates (Raja et al., 2019).

Interestingly, aberrant intermediary metabolism is also a hallmark of Barth syndrome, a human disease caused by mutations in the CL-remodeling enzyme, tafazzin (Schlame & Ren, 2006). Analyses of plasma metabolites point to alterations in carbohydrate and amino acid metabolism in Barth syndrome patients compared to healthy controls (Cade et al., 2013). Remarkably, perturbations in energy metabolism, in particular in the levels of TCA intermediates and amino acid metabolites, were recently also found in a tafazzin knock out mouse cell line (Raja & Greenberg, 2014), linking CL to energy metabolism. Surprisingly, despite these important functions, CL is dispensable for viability in *Escherichia coli, S. cerevisiae* and mammalian cell lines (Jiang, Gu, Granger, & Greenberg, 1999; Nishijima et al., 1988; Raemy et al., 2016). In contrast, CL is essential for survival of *Trypanosoma brucei* procyclic forms in culture (Serricchio & Bütikofer, 2012).

The biosynthesis of CL occurs on the matrix side of the inner mitochondrial membrane in four steps via the intermediates phosphatidic acid, CDP-diacylglycerol, phosphatidylglycerophosphate and phosphatidylglycerol (PG). In the final step, in most eukaryotes CL is formed from PG and CDP-diacylglycerol by eukaryotic-type CL synthases. In contrast, in certain parasitic protozoa and prokaryotes, CL is formed from two PG molecules (or from one PG and one phosphatidylethanolamine molecule) by prokaryotic-type CL synthases (Schlame, 2008). Most previous studies aiming to investigate the importance of CL for inner mitochondrial membrane protein complexes and mitochondrial function were carried out in cells after knocking out key enzymes in the CL biosynthetic pathway. While this approach has yielded a wealth of information on steady-state changes in mitochondrial protein levels and mitochondrial function, such systems are not suited to detect subtle changes and adaptations that occur *during* CL depletion. In *T. brucei* procyclic form CL synthase (TbCls) conditional knock-out parasites, we have recently shown that gradual CL depletion not only affected the stability of mitochondrial respiratory complexes but also decreased the levels of several mitochondrial proteins that have not been previously linked to CL (Schädeli et al., 2019). These proteins, named CL-dependent proteins (CLDPs), were identified by comparing the proteomes of procyclic form trypanosomes at different time-points after induction of CL depletion using an unbiased mass spectrometry-based approach (Schädeli, et al., 2019).

*T. brucei* parasites cycle between the fly host, Glossina *spp*., and mammals, and are the causative agents of human African trypanosomiasis and nagana in domestic animals. To cope with the fundamentally different host environments during its life cycle, the parasite has adapted its energy metabolism to the availability of different nutrients. In the tsetse fly midgut, *T. brucei* procyclic forms thrive mostly on amino acids, express functional TCA cycle enzymes and generate ATP by substrate-level and oxidative phosphorylation (Hannaert, Bringaud, Opperdoes, & Michels, 2003). In contrast, bloodstream form parasites produce cellular ATP via cytosolic substrate-level phosphorylation by aerobic glycolysis. As a result of the constant availability of glucose in the blood, bloodstream form trypanosomes minimize mitochondrial energy metabolism and down-regulate key enzymes of the TCA cycle as well as the cytochrome *c*-containing respiratory chain protein complexes (Zíková, Hampl, Paris, Tyc, & Lukes, 2016). In the absence of the proton-pumping complexes III and IV, the electrochemical potential across the inner mitochondrial membrane in *T. brucei* bloodstream forms is generated and maintained by the hydrolytic activity of the F_o_F_1_-ATP synthase (Nolan & Voorheis, 1992; Schnaufer, Clark-Walker, Steinberg, & Stuart, 2005). As a result, the mitochondrion differs morphologically and metabolically between these life cycle stages (Priest & Hajduk, 1994; Smith, Bringaud, Nolan, & Figueiredo, 2017; Tielens & van Hellemond, 2009; Vickerman, 1985; Zíková, Verner, Nenarokova, Michels, & Lukes, 2017).

In the present study, we have exploited the unusual role of the *T. brucei* bloodstream form mitochondrion to study the effects of CL depletion in a cell lacking a canonical respiratory chain and an F_o_F_1_-ATP synthase working in reverse direction compared to most other eukaryotes. We generated bloodstream form TbCls conditional knock-out parasites and examined time-dependent changes in protein levels and metabolites during CL depletion using quantitative comparative mass spectrometry. Our results show that ablation of TbCls expression causes a decreased respiration rate and rapid drop in cellular ATP levels resulting in a reduction of the mitochondrial membrane potential (ΔΨm). Proteomic and metabolomic analyses revealed a large number of proteins that were down-regulated during CL depletion and, unexpectedly, a set of mitochondrial proteins involved in energy metabolism that were up-regulated, possibly to counteract the loss of ATP and ΔΨm.

## Results

### Generation and characterization of bloodstream form conditional TbCls knock-out parasites

To study the importance of CL in an organism lacking a canonical cytochrome *c*-containing electron transport chain (ETC), we generated *T. brucei* bloodstream form TbCls conditional knock-out (TbCls KO) mutants by deleting both endogenous TbCls genes and expressing a tetracycline-dependent HA-tagged ectopic copy of TbCls (Fig. 1A). Replacement of both TbCls alleles in the conditional TbCls knock-out mutant was confirmed by Southern blotting (Fig. S1A) and the presence and correct genomic integration of the two antibiotic resistance genes was verified by PCR using gene-specific primers (Fig. S1B). Removal of tetracycline from the culture medium for 24 h resulted in a reduction of TbCls mRNA (Fig. 1B), disappearance of HA-tagged TbCls (Fig. 1C) and growth arrest after 48 h followed by parasite death (Fig. 1D), demonstrating that expression of TbCls is essential in *T. brucei* bloodstream forms and that growth of parasites can be maintained by expressing a tetracycline-dependent HA-tagged copy of TbCls. *De novo* production of CL was analyzed by *in vivo* [^3^H]-glycerol labeling of TbCls KO parasites and revealed that after TbCls depletion for 24 h no label was incorporated into CL (Fig. 1E), showing that ablation of TbCls expression inhibits CL synthesis. To examine whether CL depletion leads to defects in mitochondrial structural and functional integrity, we first analyzed mitochondria ultrastructure by transmission electron microscopy. Electron micrographs of TbCls KO parasites after 0 and 48 h of CL depletion showed no obvious morphological defects in mitochondria (Fig. 2A). In contrast, examination of the mitochondrial membrane potential ΔΨm using the potential-dependent dye MitoTracker Red and fluorescence microscopy revealed that parasites lacked the typical mitochondrial staining observed in control trypanosomes after depletion of TbCls for 24 h (Fig. 2B). In addition, quantification of the mitochondrial membrane potential in live cells with tetramethylrhodamine ethyl ester (TMRE) using flow cytometry showed a marked reduction in ΔΨm by ~50% after 24 h and ~75% after 48 h of TbCls depletion (Fig. 2C).

**Figure 1:**
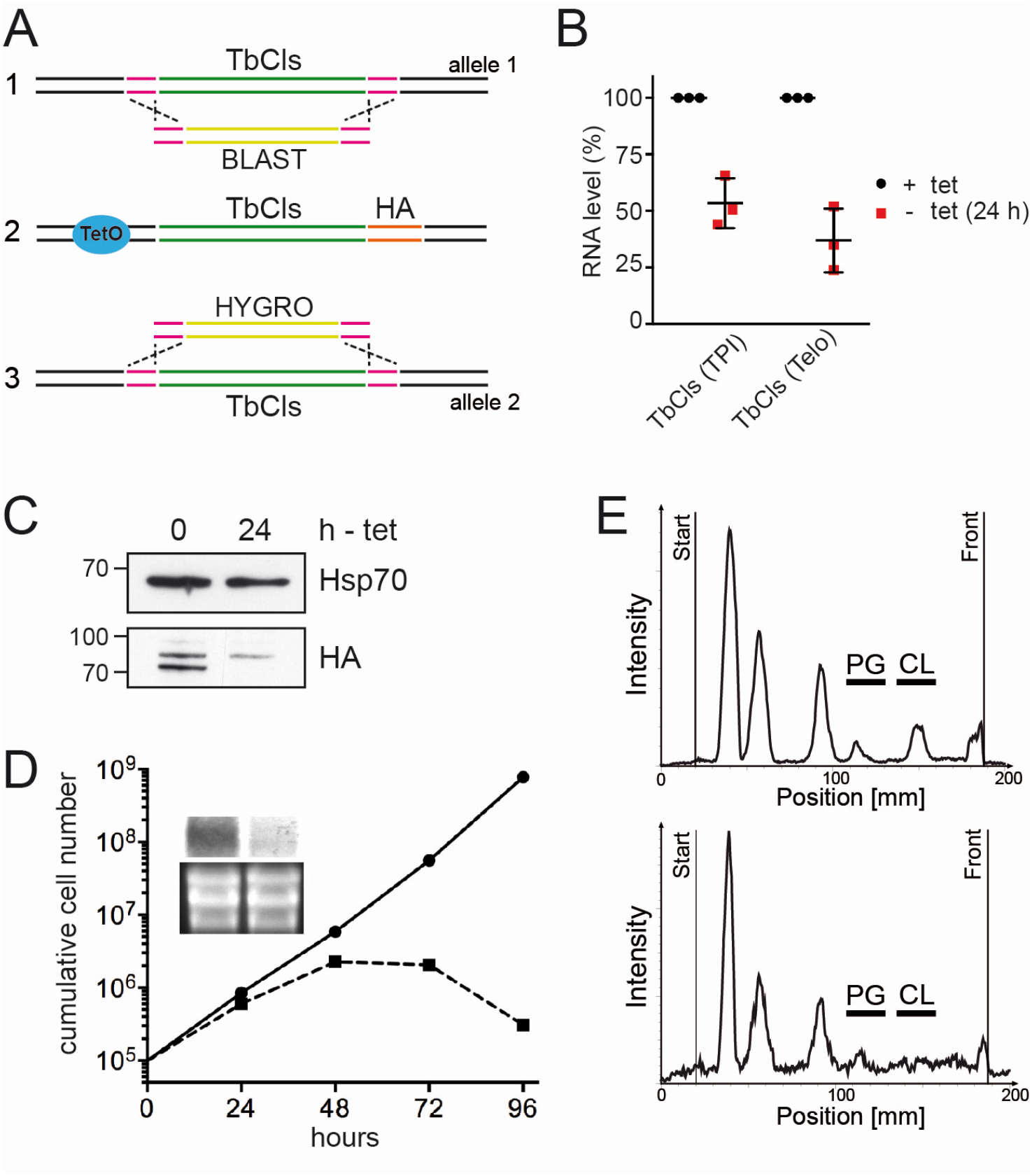
Characterization of conditional TbCls KO bloodstream form parasites. A) The strategy to generate an inducible TbCls KO involved replacement of one allele with a resistance gene conferring blasticidin resistance, introduction of an ectopic tetracycline-inducible C-terminally HA-tagged TbCls ORF followed by replacement of the second TbCls allele with a hygromycin resistance gene. B) Quantitative PCR assessment of TbCls mRNA after 24 h of tetracycline removal relative to the two housekeeping genes triosephosphate isomerase (TPI) or telomerase (Telo). C) Immunoprecipitation and immunoblot analysis of HA-tagged TbCls before (0) or after 24 h of tetracycline removal. Hsp70 was used as input control. D) Growth curve of conditional TbCls KO cells cultured in the presence (filled circles) or absence (filled squares) of tetracycline to induce TbCls depletion. E) *In vivo* metabolic labeling of TbCls KO parasites before (top panel) or after TbCls depletion for 24 h (bottom panel) with [^3^H]-glycerol for 6 h followed by phospholipid extraction and analysis using thin-layer chromatography and radioisotope scanning.

**Figure 2:**
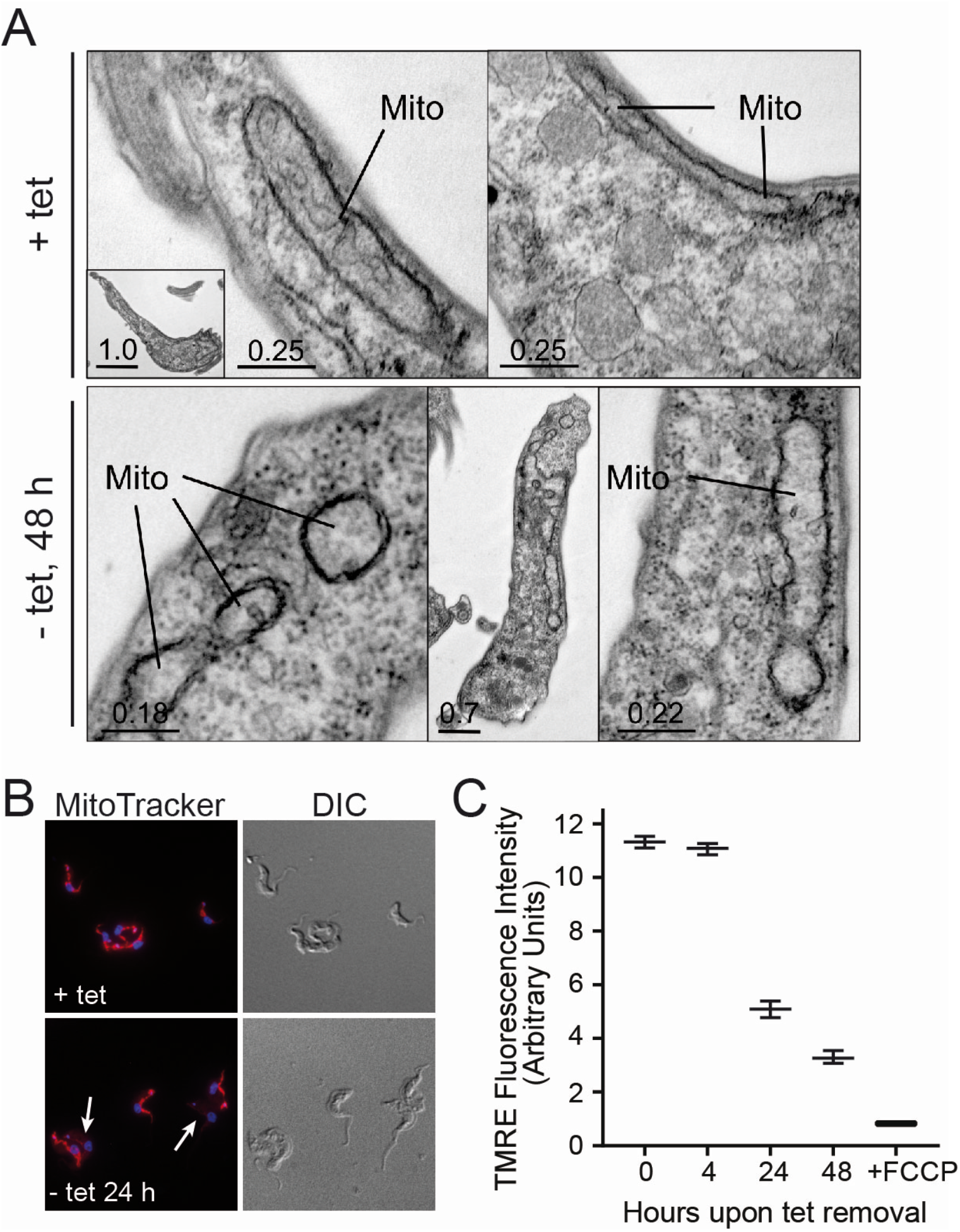
Mitochondrial characterization of TbCls KO parasites. A) Transmission electron microscopy of TbCls KO cells cultured in presence (+ tet) or absence (- tet) of TbCls expression for 48 h. B) MitoTracker staining of live parasites before (+tet) or after (-tet) TbCls depletion for 24 h. Arrows point to cells lacking MitoTracker staining. C) The ΔΨm of TMRE-stained TbCls KO cells before (0) or after depletion of TbCls for 4, 24 or 48 h was measured using flow cytometry. FCCP was used to dissipate ΔΨm. The median fluorescence ± S. D. from three biological replicates is depicted.

**Figure S1:**
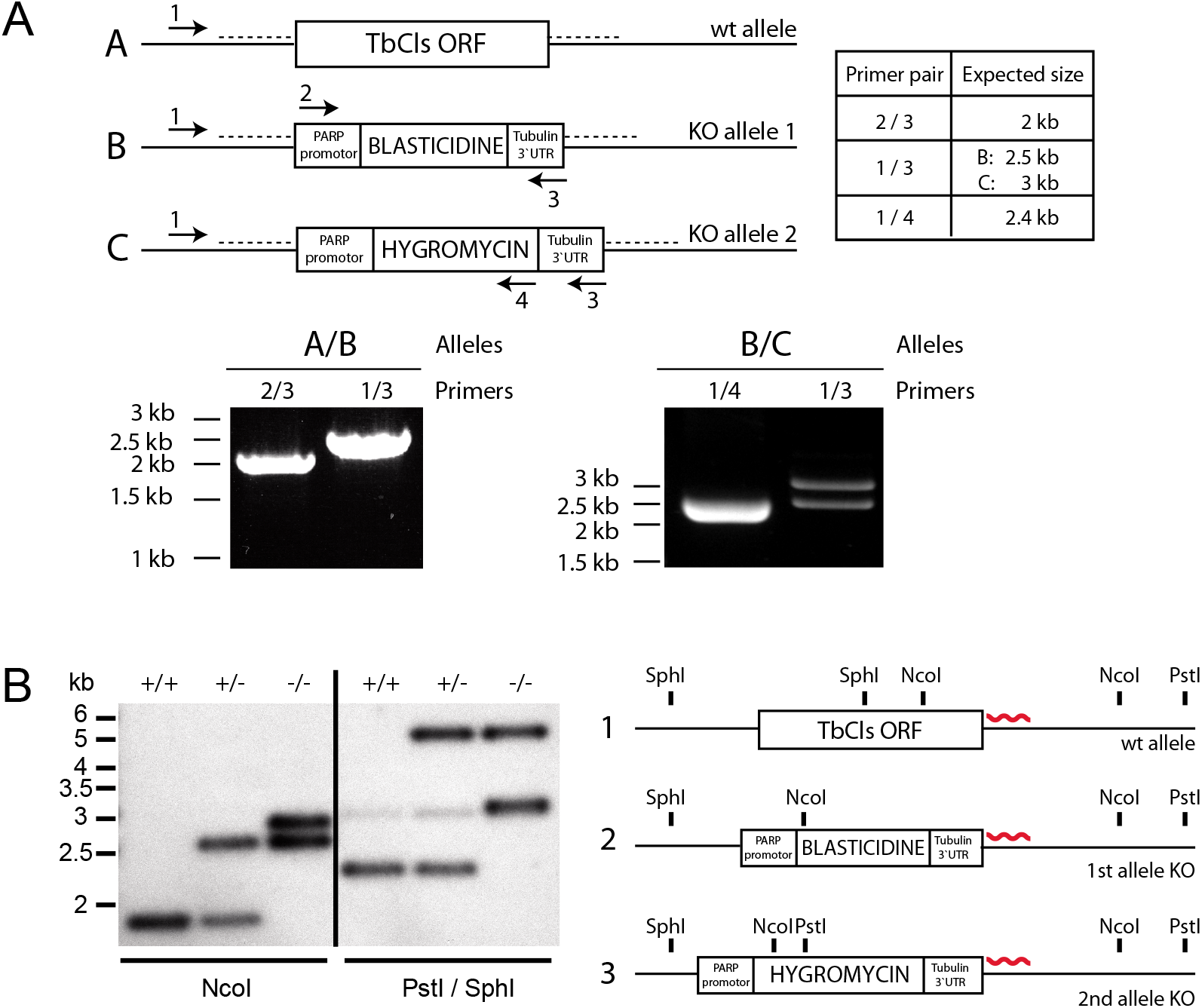
Validation of TbCls KO parasites. A) PCR verification of single-allele (alleles A/B) and the final double allele knockout cell line (alleles B/C). Sizes of expected PCR fragments and primers used for amplification are shown. B) Northern blot analysis of wt (+/+), single-allele (+/-) and double-allele TbCls KO (-/-) after digestion of genomic DNA with either NcoI or PstI/SphI. Expected restriction sites are depicted in the schematic, and the binding site of the hybridizing probe is shown as a red wavy line.

### F_o_F_1_-ATPase complex organization and activity during CL depletion

In *T. brucei* bloodstream forms, the ΔΨm is maintained by ATP-dependent proton pumping activity of F_o_F_1_-ATPase complexes (Nolan & Voorheis, 1992; Schnaufer, et al., 2005). To test if the drop in ΔΨm was caused by destabilization of F_o_F_1_-ATPase complexes as a result of decreased CL levels, light blue native gel electrophoresis was performed. Native complexes were detected after immunoblotting with specific antibodies against F_1_ subunits β and p18 or F_o_ subunit Tb2 (Gahura et al., 2018; Subrtova, Panicucci, & Zíková, 2015). The results show a mild reduction in the abundance of the monomeric/dimeric state of the F_o_F_1_-ATPase complex accompanied by an accumulation of the F_1_ assembly intermediate after 24 h of TbCls depletion, which becomes more pronounced after 48 h of TbCls depletion (Fig. 3A). To assess the ability of F_o_F_1_-ATPase complexes to pump protons, we measured uptake of safranin O into mitochondria of digitonin-permeabilized cells in the presence of exogenously added ATP. Safranin O is a lipophilic cationic dye that, upon membrane potential-dependent uptake into mitochondria, undergoes a spectral change and fluorescence quenching that can be used to estimate ΔΨm (Figueira, Melo, Vercesi, & Castilho, 2012). Relative to control parasites expressing TbCls, we observed a ~20% reduction of safranin O uptake after depletion of TbCls for 24 h, while a decrease of ~40% was detected after 48 h of TbCls ablation (Fig. 3B, C, D). The difference between the ΔΨm measured by TMRE in intact cells (see Fig. 2C) and safranin O uptake measured in digitonin-permeabilized cells in the presence of excess ATP suggests that early during depletion of TbCls, i.e. after 24 h, lack of ATP rather than loss of F_o_F_1_-ATPase function is responsible for the drop in ΔΨm. Indeed, quantification of the ADP/ATP ratio and total ATP levels after 24 h of TbCls depletion revealed an increase in the ADP/ATP ratio (Fig. 3E) and a drop in cellular ATP (Fig. 3F) compared to control cells. In summary, after 24 h of TbCls depletion, F_o_F_1_-ATPase structure and activity appear to be only marginally affected, yet the ΔΨm is strongly reduced, likely due to the drop in cellular ATP.

**Figure 3:**
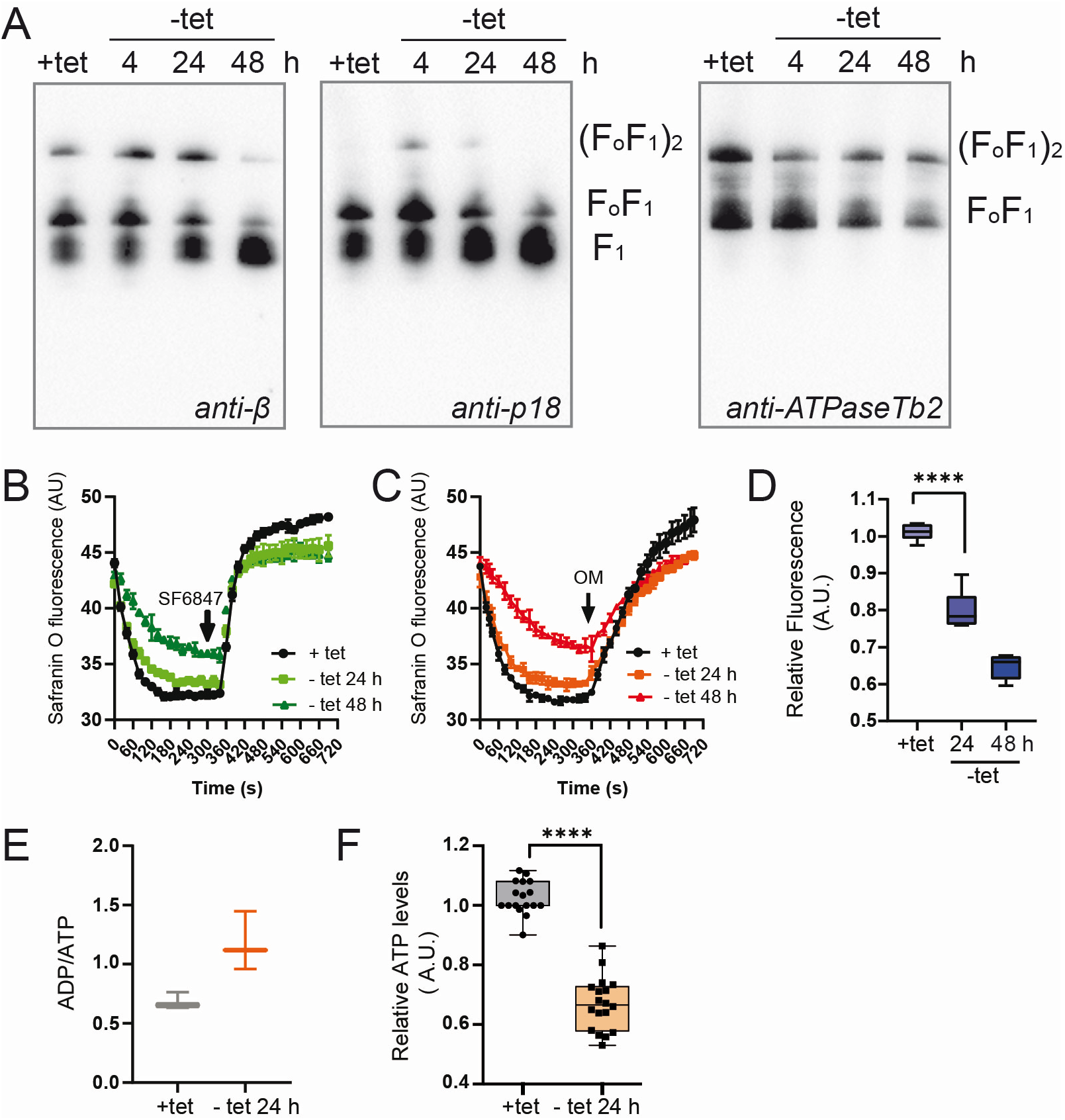
Functional assessment of the mitochondrial F_o_F_1_-ATP synthase complex. A) Native F_1_- and F_o_F_1_-ATPase complexes were visualized using light blue native electrophoresis. Purified mitochondria from +/-tet cultures were lysed with dodecyl maltoside, fractionated on 3-12% Bis-Tris gels and blotted onto a PVDF membrane. The F_1_-ATPase and F_o_F_1_-ATPase monomer and dimers were detected using polyclonal antibodies recognizing F_1_-ATPase subunits β and p18, or F_o_F_1_-ATPase subunit Tb2. B-C) The ability of TbCls KO parasites cultured in the presence (+tet) or absence of tetracycline (-tet) for 24 or 48 h to establish ΔΨm *in vitro* was measured using a fluorescent indicator, safranin O, in the presence of excess ATP. The reaction was triggered by addition of digitonin. The extent of the spectral change correlates linearly to ΔΨm. The ΔΨm was dissipated after the addition of the uncoupler SF6847 (B) or oligomycin (C). D) The fluorescence changes between the time points 0 and 340 s were normalized to the value of TbCls KO (+tet) and plotted using Graph Pad Prism 8.2.1 (n=3); **** p < 0.0001, Student’s t-test). E,F) Comparison of ADP/ATP ratios (E) and cellular ATP content (F) in TbCls KO parasites cultured in the presence (+tet) or absence of tetracycline (-tet) for 24 h (mean values ± S.D., n>3; **** p < 0.0001).

### Metabolomic analyses of TbCls-depleted trypanosomes

To identify possible metabolic changes caused by TbCls depletion, we performed metabolomic analyses of parasites after 12 h and 24 h of TbCls depletion and compared metabolite levels to control cells. Interestingly, of a total of 5050 detected peaks, we observed significant changes (p-values <0.05) in several key metabolites associated with energy deprivation. While the levels of phosphoarginine, acetoacetate and oxidized hexoses were decreased, AMP levels were increased (Table 1). Phosphoarginine represents the equivalent to phosphocreatine in vertebrates by providing high-energy phosphate groups to replenish ATP levels on a short time-scale and is produced by multiple isoforms of phosphoarginine kinases (Voncken, Gao, Wadforth, Harley, & Colasante, 2013). Acetoacetate represents a ketone body produced under conditions of starvation and has been identified in *T. brucei* before (Shah, Hickey, Capasso, & Palenchar, 2011), whereas oxidized hexoses, e.g. D-gluconate, are substrates for the pentose-phosphate pathway for nucleotide biosynthesis and formation of NADPH, a key reducing agent for protection against oxidative stress in trypanosomes (Kovarova & Barrett, 2016).

**Table1:**
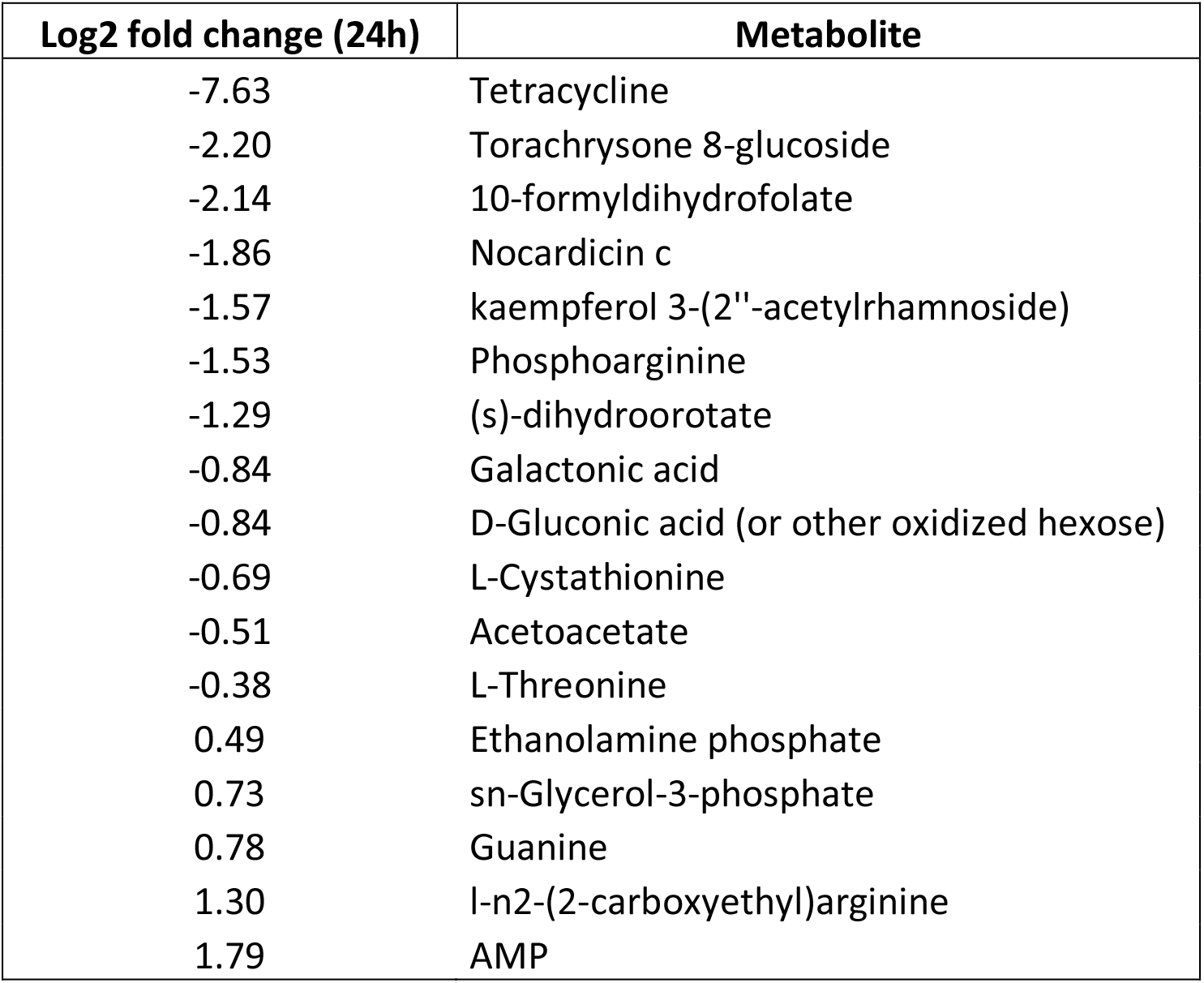
Metabolomic changes triggered by depletion of TbCls. List of mass spectrometry signals matched to known standards. The log2 fold change between metabolites found in parasites expressing TbCls and depleted of TbCls for 24 h are listed. Metabolites shown had a significance at a p-value of <0.05.

Pathway analyses using Polyomics integrated Metabolomics Pipeline PiMP (Gloaguen et al., 2017) revealed no consistent changes in metabolite levels in the glycolytic pathway, the TCA cycle or amino acid metabolism, while glycerolipid metabolism was the most significantly changed pathway between control and 24 h TbCls-depleted parasites, highlighted by increased levels of glycerol and glycerol-3-phosphate (Fig. S2). Together, the metabolomic analyses support the above observations that parasites after 24 h of TbCls depletion are in an energy-deprived state.

**Figure S2:**
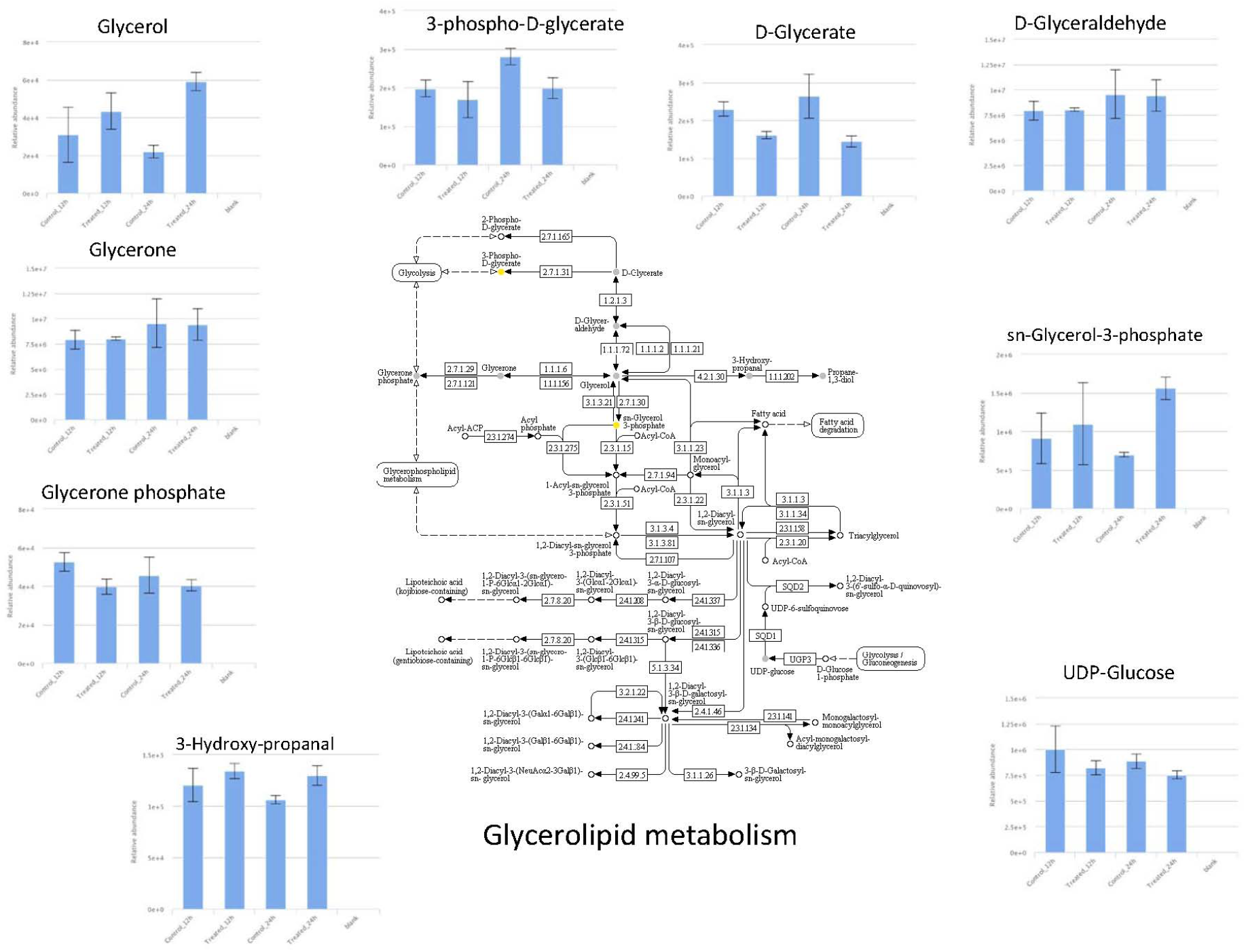
Changes of metabolite levels of glycerolipid metabolism. The blue bars depict metabolite levels of control (+ tet, 0h), TbCls depleted (-tet, 12h), control (+ tet, 24 h) and TbCls depleted (-tet, 24 h) parasites, respectively.

### Proteomic changes during CL depletion

Using stable isotope labeling with amino acids in cell culture (SILAC) and mass spectrometry we subsequently compared the proteomes of mitochondria-enriched extracts from *T. brucei* bloodstream forms after depletion of TbCls for 12 h and 24 h with parasites expressing TbCls. Previous experiments have shown that labeled amino acids are uniformly incorporated into the proteome in *T. brucei* parasites (Cirovic & Ochsenreiter, 2014; Schädeli, et al., 2019) and that parasite growth was not affected by the different culture media used for SILAC (Fig. S3).

**Figure S3:**
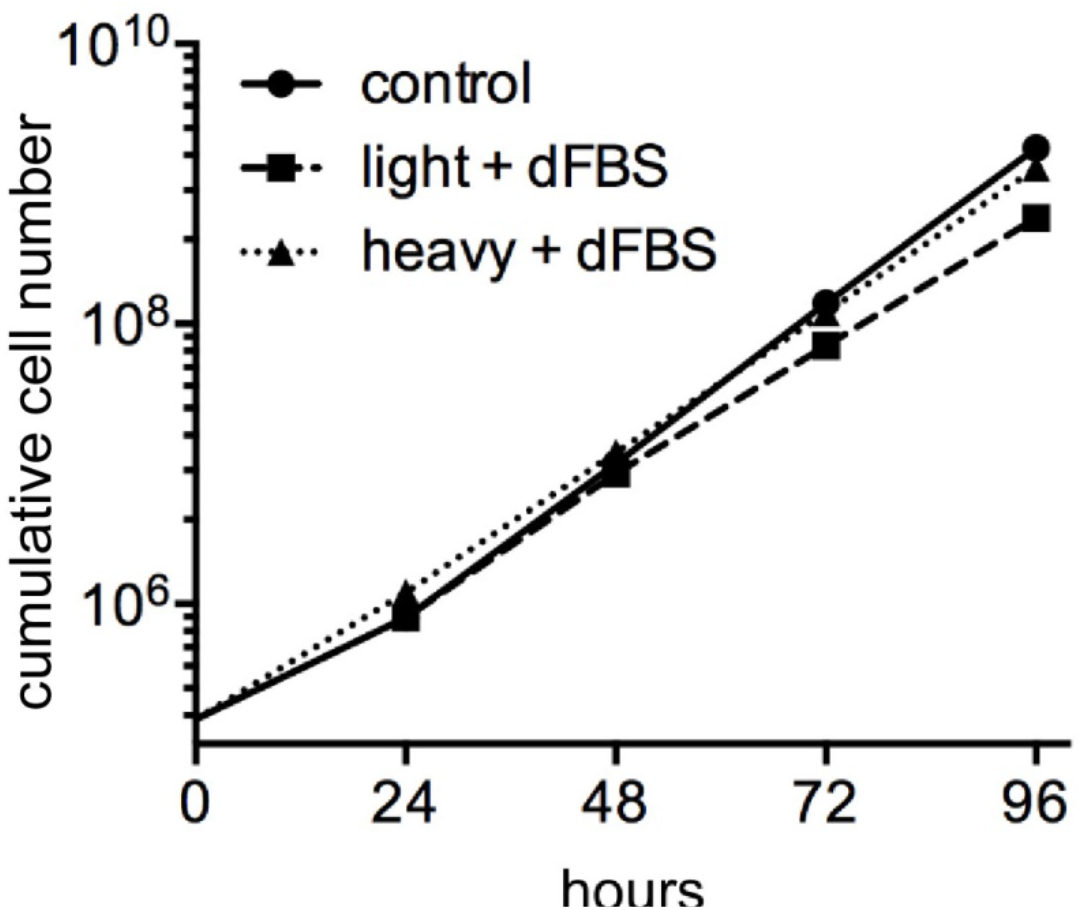
Growth of TbCls KO parasites cultured in light and heavy medium containing normal or isotope-labeled amino acids. Parasites were cultured in medium containing L-arginine and L-lysine and dialyzed fetal bovine serum (light + dFBS) or isotope labeled L-arginine and L-lysine with dialyzed fetal bovine serum (heavy + dFBS). Control parasites were cultured in light medium with non-dialyzed fetal bovine serum (control).

Our analyses revealed >1100 proteins in each of the biological triplicates from all three time points (0 h, 12 h and 24 h of TbCls ablation). We identified a large number of proteins with altered expression levels after 12 h and 24 h of TbCls depletion compared to control trypanosomes (Fig. 4A-C), with >110 proteins with fold-changes of down-regulation of ≥1.6 (Fig. 4D, Table S1) and >35 proteins with fold-changes of up-regulation of ≥1.4 (Fig. 4D, Table S2).

**Figure 4:**
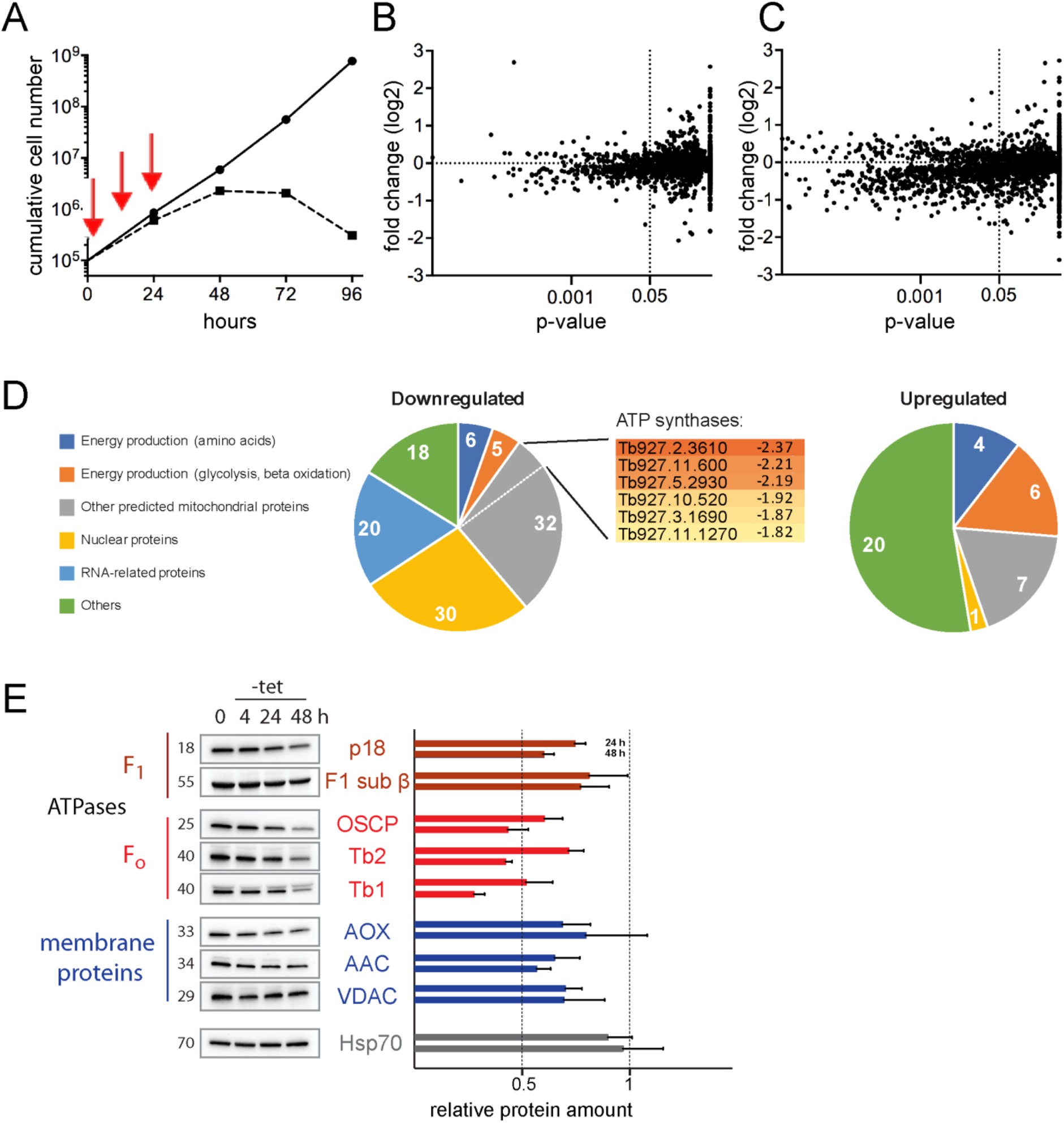
Cardiolipin-induced changes in protein levels. A-C) SILAC analysis of protein abundance during depletion of TbCls. Membrane-enriched fractions from TbClsKO parasites cultured for 0 h, 12 h or 24 h (indicated by red arrows in A) in the absence of tetracycline to induce ablation of TbCls expression were analyzed by mass spectrometry. The p-value of each data point was determined by a two-sample t-test between time points 0 h and 12 h (B) and 0 h and 24 h (C) from three biological independent experiments and was plotted against the fold change in protein abundance after 12 h and 24 h, respectively, relative to the control sample taken at time 0 h. D) Grouping of proteins that were down- or up-regulated upon TbCls depletion for 24 h. Gene IDs for down-regulated ATP synthase subunits are shown. E) Immunoblot analyses of TbCls KO cells grown in the presence (0) or absence of tetracycline for 4, 24 and 48 h. Signals from -tet samples are shown relative to control samples (n ≥ 3).

Among the down-regulated proteins, (predicted) mitochondrial proteins comprised the largest group (Fig. 4D, Table S1). Interestingly, we identified two known subunits of ATP synthase, Tb1 (Tb927.10.520) and Tb2 (Tb927.5.2930), and several proteins annotated as novel subunits of ATP synthase (Tb927.2.3610, Tb927.11.600, Tb927.3.1690, Tb927.11.1270) (Fig. 4D, Table S1). In addition, among the down-regulated proteins, we identified six (putative) proteins involved in amino acid metabolism, with two of them representing putative amino acid transporters of the AAT4 family with unknown specificities and localization (Tb927.4.4830/4850/4870; Tb927.4.4860), one being a putative mitochondrial amino acid transporter of the AAT17 family (Tb927.11.15950) and the other annotated as putative lysine transport protein (Tb927.11.15840/15860). Furthermore, down-regulation was also observed for several mitochondrial and cytosolic proteins involved in energy metabolism, such as putative enoyl-CoA hydratase (Tb927.11.16480), malic enzyme (Tb927.11.5450), putative hydroxymethylglutaryl-CoA lyase (Tb927.4.2700), putative acyl-CoA synthetase (Tb927.6.2010), putative NADH-cytochrome b5 reductase (Tb927.5.1470), putative adenylate kinase (Tb927.10.2540) and pyruvate kinase 1 (Tb927.10.14140). The list of down-regulated proteins also comprised proteins involved in nucleotide metabolism (two adenosine transporters, ribonucleoside-diphosphate reductase) and *N*-glycosylation (putative dolichyl-P-Man:GDP-Man7GlcNAc2-PP-dolichyl alpha-1,6-mannosyltransferase, putative UDP-Gal/UDP-GlcNAc-dependent glycosyltransferase) and two large sets with nuclear/nucleolar proteins and proteins involved in RNA synthesis/processing (Table S1).

Among the up-regulated proteins, we identified proteins involved in energy production via carbohydrate metabolism, including the bloodstream form-specific glucose transporter THT1 (Tb927.10.8440), hexokinase (Tb927.10.2010), fructose-bisphosphate aldolase (Tb927.10.5620) and glyceraldehyde 3-phosphate dehydrogenase (Tb927.10.6880) (Fig. 4D, Table S2), and several mitochondrial enzymes connecting amino acid metabolism to the TCA cycle, such as glutamate dehydrogenase (Tb927.9.5900), putative hydroxyglutarate dehydrogenase (Tb927.10.9360), putative 2-oxoglutarate dehydrogenase, component E1 (Tb927.11.9980) and succinate dehydrogenase assembly factor 2 (Tb927.6.2510). Remarkably, several nutrient and ion transporters ranked among the most highly up-regulated proteins, such as putative mitochondrial amino acid transporter AAT7 (Tb927.8.7610/8.7640; 2.3-fold up-regulation), aquaglyceroporin 1 (Tb927.6.1520; 2.1-fold up-regulation), mitochondrial folate transporter (Tb927.8.3650; 1.9-fold up-regulation) and putative mitochondrial V-type ATPase (Tb927.4.1080; 1.5-fold up-regulation).

To verify CL-dependent down-regulation of selected mitochondrial proteins, protein extracts from *T. brucei* bloodstream forms after down-regulation of TbCls were analyzed by SDS-PAGE and immunoblotting using antibodies recognizing F_o_F_1_-ATPase subunits, mitochondrial membrane proteins or mitochondrial matrix proteins (Fig. 4E). The results after TbCls depletion revealed decreased levels of several F_o_ subunits (>20 and >50% reduction after 24 h and 48 h, respectively) and F_1_ subunits (>20% reduction after 24 h) of the F_o_F_1_-ATPase complex (Fig. 4E, top two panels) and the inner mitochondrial membrane proteins alternative oxidase (AOX; Tb927.10.7090; ~25% reduction after 24 h) and ADP/ATP carrier (AAC; Tb927.10.14830; ~30% reduction after 24 h) (Fig. 4E, middle panel). These results are in line with the SILAC/mass spectrometry data showing reduced levels of several ATP synthase subunits (see Table S1) and a reduction in AOX (~20% decrease after 24 h; AAC was not detected in the SILAC experiments).

In bloodstream form trypanosomes, glycolysis and thus ATP production is directly coupled to mitochondrial respiration via a truncated ETC composed of glycerol-3-phosphate dehydrogenase and alternative oxidase (Opperdoes, Borst, Bakker, & Leene, 1977). To assess if the CL-induced drop in ATP levels is linked to decreased cellular respiration, we measured oxygen consumption rates of live parasites during the depletion of TbCls. Our results show that O_2_ consumption after 24 h and 48 h of CL depletion was significantly reduced relative to control cells (Fig. 5A, B). Since AOX is a dimer bound to the inner mitochondrial membrane via a hydrophobic region in an interfacial fashion (Shiba et al., 2013), we tested if CL may be involved in membrane binding of AOX. We performed carbonate extraction experiments with isolated mitochondria from parasites before and after depletion of CL and found that AOX expression is indeed decreased (Fig. 5C; see also Fig. 4E), however, partitioning of AOX between pellet and supernatant fractions was unchanged between TbCls-depleted and control parasites (Fig. 5C), demonstrating that depletion of CL affected AOX steady-state levels but not binding to the inner mitochondrial membrane.

**Figure 5:**
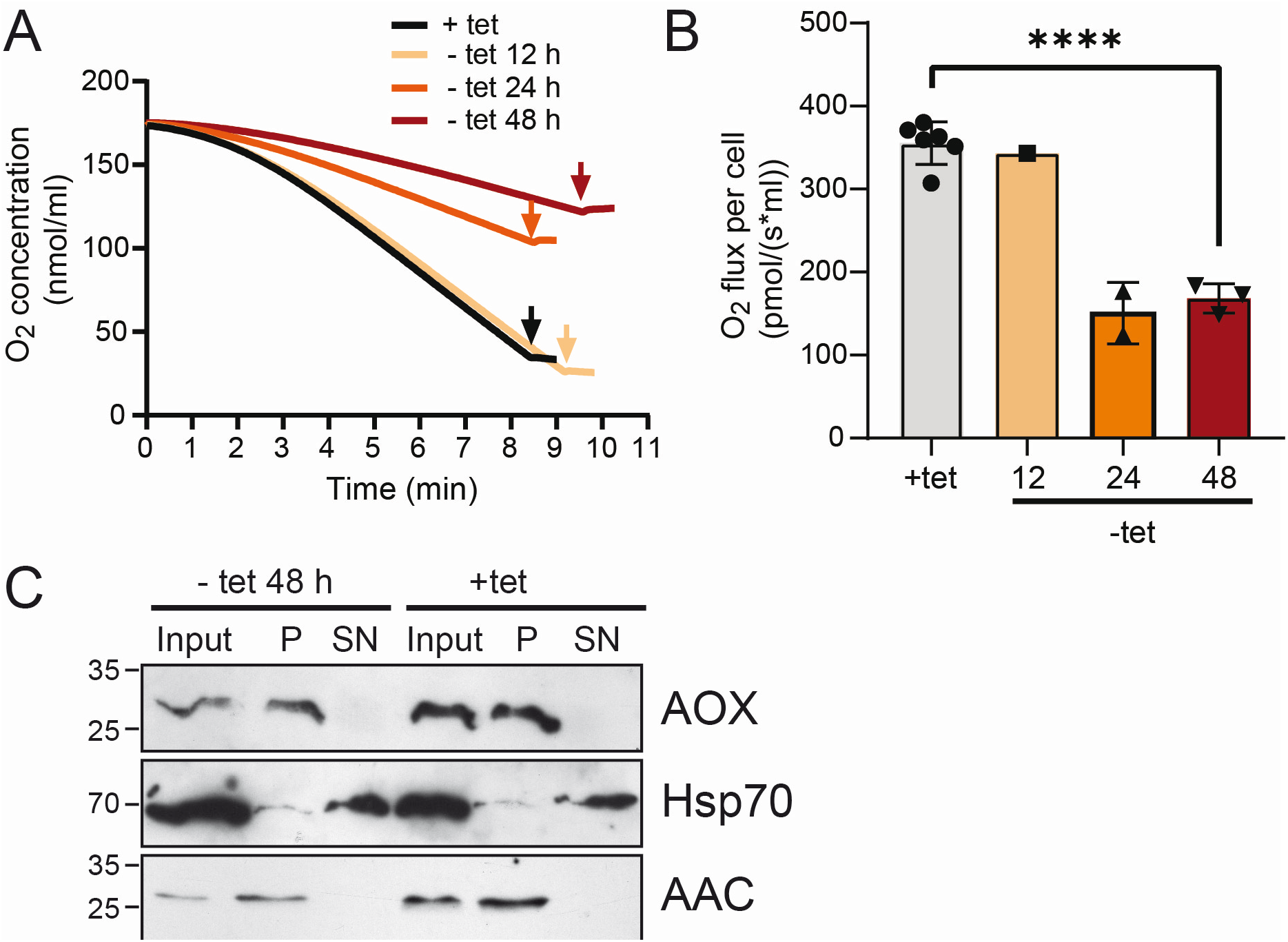
Oxygen consumption is affected by CL depletion. A) Representative respiratory traces of TbCls KO parasites cultured in the presence (+tet) or absence of tetracycline (-tet) for 24 or 48 h were determined using high-resolution respirometry (Oroboros Instrument). The graphs depict changes in oxygen concentration (nmol/ml) in the media over time (x-axes) (left panel). Arrows indicate time of addition of the inhibitor SHAM. B) Oxygen flow in TbCls KO parasites cultured in the presence (+tet) or absence of tetracycline (-tet) for 24 or 48 h is expressed as respiration per million cells (pmol x s^−1^ x 10^−6^ cells) (right panel). C) Carbonate extraction from mitochondria-enriched fractions of TbCls KO parasites grown in presence or absence of TbCls expression for 48 h. Fractionation of AOX was compared to the soluble protein Hsp70 and the integral membrane protein AAC.

## Discussion

High-resolution structures have revealed specific binding of CL to mammalian and yeast respiratory complexes I (Fiedorczuk, et al., 2016), III (Lange, et al., 2001; Solmaz & Hunte, 2008) and IV (Shinzawa-Itoh, et al., 2007) of the inner mitochondrial membrane. In addition, CL stimulates the activities of these complexes and stabilizes super-complex formation (Mileykovskaya & Dowhan, 2014). As a result, in CL-deficient mammalian, yeast and plant cells, mitochondrial ultrastructure and function, in particular respiration, is affected. In spite of these defects, the cells are viable (Jiang, Rizavi, & Greenberg, 1997; Pineau et al., 2013; Raemy, et al., 2016). In contrast, CL is essential for the survival of *T. brucei* procyclic forms in culture (Serricchio & Bütikofer, 2012, 2013). Here, we investigated the effects of CL depletion in *T. brucei* bloodstream forms, i.e. in cells lacking a functional ETC and ATP production via oxidative phosphorylation (Hannaert, et al., 2003).

To control *de novo* CL synthesis, we deleted both alleles of TbCls in *T. brucei* bloodstream forms and introduced a tetracycline-dependent copy of HA-tagged TbCls. Upon removal of tetracycline from the culture medium, TbCls expression was down-regulated and *de novo* synthesis of CL was inhibited. These conditions resulted in growth arrest of parasites followed by cell death, demonstrating that the production of CL in *T. brucei* bloodstreams forms is essential for parasite survival in culture.

Interestingly, after inhibition of CL synthesis, proteomic analyses revealed markedly reduced levels of many mitochondrial proteins. Most notably, several confirmed and predicted subunits of the inner mitochondrial membrane F_o_F_1_-ATP synthase were severely down-regulated. CL is tightly associated with F_o_F_1_-ATP synthase, (Eble, Coleman, Hantgan, & Cunningham, 1990; Muhleip, McComas, & Amunts, 2019), and crystal structures have revealed CL binding sites in the rotor-stator interface, the dimer interface, and in a peripheral F_o_ cavity (Muhleip, et al., 2019). Molecular modeling suggests that CL participates in proton translocation through the membrane domain (Duncan, Robinson, & Walker, 2016). Upon CL depletion in *Drosophila* flight-muscle mitochondria, F_o_F_1_-ATP dimers are destabilized and their lateral membrane distribution is distorted (Acehan, et al., 2011). In *T. brucei* bloodstream forms, we observed destabilization of F_o_F_1_-ATP dimers as well as monomers during CL depletion leading to accumulation of F_1_-ATPase subcomplex. Uncoupling of the F_o_ and F_1_ parts of ATP synthase has been observed before in naturally occurring and drug-induced dyskinetoplastic bloodstream form trypanosomes (Schnaufer, Domingo, & Stuart, 2002), which maintain the mitochondrial membrane potential in an F_o_-independent fashion (Schnaufer, et al., 2005) by coupling F_1_-ATPase activity with electrogenic exchange of the cytosolic ATP^4-^ for the mitochondrial matrix ADP^3-^ by the AAC. However, this mechanism is conditioned by acquiring specific mutations in the F_1_ subunits (Dean, Gould, Dewar, & Schnaufer, 2013; Lai, Hashimi, Lun, Ayala, & Lukes, 2008), which presumably increase the enzyme’s capacity to hydrolyze ATP. Without these mutations, bloodstream form cells are not capable of maintaining mitochondrial membrane potential in an F_o_-independent mode (Schnaufer, et al., 2005). These observations are consistent with our results of the ΔΨm measurements *in vivo* by TMRE and *in vitro* by safranin O at 48 h of CL depletion. The decreased levels of the coupled F_o_F_1_-ATPase led to a substantial drop in ΔΨm. Interestingly, at 24 h of TbCls depletion, the ΔΨm measured *in vivo* by TMRE was already decreased by 50%, while *in vitro* safranin O uptake measurements in presence of excess ATP revealed only a 20% reduction in ΔΨm, suggesting that at this early timepoint it is the lack of the substrate, i.e. ATP, rather than the dysfunctional F_o_F_1_-ATPase that leads to the observed ΔΨm phenotype.

In addition to a down-regulation of F_o_F_1_-ATP synthase subunits, CL depletion resulted in a decrease of several (putative) transporters. It has been shown before that CL associates with and affects the activity of the mitochondrial AAC (Beyer & Klingenberg, 1985; Claypool, Oktay, Boontheung, Loo, & Koehler, 2008). In addition, CL stimulates the activities of the carnitine/acylcarnitine transporter (Paradies, Ruggiero, Petrosillo, & Quagliariello, 1997) and the phosphate carrier (Kadenbach, Mende, Kolbe, Stipani, & Palmieri, 1982), and it is possible that additional mitochondrial carriers also depend on CL for proper function (Claypool, 2009). Although the subcellular localization of several of the *T. brucei* transporters down-regulated after CL depletion has not been experimentally established, our data suggest that they may localize to the inner mitochondrial membrane, where they interact with CL and become depleted after inhibition of *de novo* CL synthesis. Two mitochondrial amino acid transporters, amino acid transporter AAT17 (Tb927.11.15950) and L-lysin transport protein (Tb927.11.15840), which have not yet been associated with CL, were down-regulated as well. Whether or not they bind CL and are down-regulated as a direct consequence of CL depletion is not clear, but deserves further attention. Similarly, although the association of AOX with the inner mitochondrial membrane was not disrupted during CL depletion, its expression was decreased. It is conceivable that the observed decrease in oxygen consumption by AOX is a direct consequence of reduced CL levels. The *T. brucei* bloodstream form cellular respiration is coupled to the glycerol-3-phosphate/dihydroxyacetone phosphate shuttle in which the mitochondrial FAD-linked glycerol-3-phosphate dehydrogenase oxidizes glycerol-3-phosphate to dihydroxyacetone phosphate and passes electrons to ubiquionone (Opperdoes, et al., 1977). The reduced ubiquinone is then oxidized by AOX. Mitochondrial glycerol-3-phosphate dehydrogenase is tightly bound to the inner mitochondrial membrane and was proposed to function in a cardiolipin-dependent fashion (Beleznai & Jancsik, 1989). It is possible that in the absence of CL, *T. brucei* glycerol-3-phosphate dehydrogenase becomes dysfunctional, and glycerol-3-phosphate is instead converted to glycerol by glycerol kinase in glycosomes, thereby resembling anaerobic conditions during which only 1 molecule of ATP is produced per 1 molecule of glucose (Bakker, Michels, Opperdoes, & Westerhoff, 1997). Indeed, the ablation of TbCls expression led to decreased ATP levels. Further, we also observed an accumulation of glycerol-3-phosphate and glycerol, indicating an improper function of glycerol-3-phosphate dehydrogenase. Interestingly, accumulation of glycerol is known to be toxic for *T. brucei* parasites, which may explain the observed increased expression of aquaglyceroporins, the transporters mediating glycerol efflux (Uzcategui et al., 2004), during CL depletion.

In addition, ablation of TbCls expression resulted in down-regulation of several mitochondrial and cytosolic enzymes. Since none of them is predicted to contain a transmembrane domain, we believe that their decrease was not a direct effect of CL depletion but a result of metabolic alterations (see below). The two sets of proteins comprising down-regulated nuclear/nucleolar proteins and proteins involved in RNA synthesis/processing likely reflect the parasite’s reduced metabolism and slowed growth after prolonged CL depletion. Together, these observations indicate that the primary effect of CL depletion in *T. brucei* bloodstream forms is on cellular respiration and the structural organization and activity of the F_o_F_1_-ATP synthase, leading to a progressive loss of the mitochondrial membrane potential and a dramatic increase in the ADP/ATP ratio as well as steady-state levels of several (mitochondrial) transporters.

Unexpectedly, and in contrast to previous findings in procyclic forms (Schädeli, et al., 2019), depletion of CL in *T. brucei* bloodstream forms led to an increase in the levels of a large number of proteins (Table S2). Many of these proteins are involved in metabolic reactions and pathways to increase ATP production. Increases in glutamate dehydrogenase (Tb927.9.5900), (putative) hydroxyglutarate dehydrogenase (Tb927.10.9360) and 2-oxoglutarate dehydrogenase (Tb927.11.9980), as well as in (putative) amino acid transporter AAT7 (Tb927.8.7610), are consistent with up-regulation of amino acid metabolism, in particular glutamate catabolism. As reported before, *T. brucei* bloodstream forms consume large quantities of glutamine, primarily as an amino group donor (Creek et al., 2013; Kim, Achcar, Breitling, Burgess, & Barrett, 2015). Recent intracellular metabolome analyses using ^13^C-labeled tracking confirmed that glutamine is catabolized to glutamate and on to 2-oxoglutarate and succinate (Johnston et al., 2019). Among the different possible pathways leading to production of succinate from glutamate (Johnston, et al., 2019), our proteomic data suggest that – during CL depletion – glutamate dehydrogenase (1.6-fold up-regulated) converts glutamate to 2-oxoglutarate which is then metabolized to succinate by 2-oxoglutarate dehydrogenase (1.74-fold up-regulated) and succinyl-CoA synthetase (Tb927.3.2230, 1.12-fold up-regulated) to produce ATP. In addition, 2-oxoglutarate can be converted to hydroxyglutarate by promiscuous action of malate dehydrogenase (Intlekofer et al., 2017) (Tb927.10.2560, 1.23-fold up-regulated) and back to 2-oxoglutarate by hydroxyglutarate dehydrogenase (1.74-fold up-regulated). Although the substrate specificity of (putative) amino acid transporter AAT7 (2.3-fold up-regulated) is currently unknown, we propose that it may be responsible for the increased uptake of glutamine into the mitochondrion of bloodstream form trypanosomes. Alternatively, increased production of glutamate could also result from AAT7-mediated uptake of proline with subsequent degradation by proline dehydrogenase (Tb927.7.210; not detected in our study) and Δ1-pyrroline-5-carboxylate dehydrogenase (Tb927.10.3210; not detected in our study).

Furthermore, our data revealed increased levels of additional enzymes and transporters involved in energy production in *T. brucei* (Table S2). Up-regulation of the glycolytic enzymes hexokinase (Tb927.10.2010), glyceraldehyde-3-phosphate dehydrogenase (Tb927.10.6880) and fructose-bisphosphate aldolase (Tb927.10.5620), and of hexose transporter THT1 (Tb927.10.8440), are again consistent with the parasite’s metabolic response to counteract CL-induced depletion of ATP. The severity of energy depletion is further reflected in a decrease in phosphoarginine levels and an increase in the arginine metabolite N2-(2-carboxyethyl)arginine. In a recent report, it was shown that in *T. brucei* bloodstream forms phosphoarginine is exclusively generated by the action of arginine kinases (Johnston, et al., 2019). Although arginine kinase knock-out parasites were viable in culture (Johnston, et al., 2019), it is believed that phosphoarginine plays a central role in regenerating ATP from ADP in situations of high (short term) energy demand (Pereira, Alonso, Torres, & Flawia, 2002).

Remarkably, while the analysis of the proteome during CL depletion revealed a pronounced decrease in the levels of several subunits of the inner mitochondrial membrane F_o_F_1_-ATP synthase (Table S1; see also above), one protein annotated as putative ATP synthase subunit (Tb927.11.9420, 1.4-fold up-regulated) was found to be increased (Table S2). Interestingly, the protein has been described as a subunit of the peripheral stalk of the vacuolar H^+^-ATPase (Huang et al., 2014). *T. brucei* bloodstream forms use V-type ATPases and mitochondrial ATPases to generate proton gradients (Schnaufer, et al., 2005), and recently functional interdependency between both types of ATPases has been reported (Baker et al., 2015). Knock-downs of V-type ATPase subunits were shown to induce kinetoplast independence and resistance of trypanosomes to isometamidium, a DNA intercalating drug that accumulates in the kinetoplast, indicating that loss of V-type ATPases affects F_o_F_1_-ATP synthase sector coupling (Baker, et al., 2015). The function of the up-regulated V-type ATPase subunit during CL depletion is not known. The observation that depletion of V-type ATPase allows for kinetoplast-independent growth (Baker, et al., 2015) suggests, however, that it’s levels were increased in response to the observed CL-dependent F_o_F_1_-ATP synthase sector uncoupling.

## Methods

Unless otherwise stated, reagents were purchased from Sigma (Buchs, Switzerland) or Merck (Darmstadt, Germany). Restriction enzymes were from Thermo Scientific (Reinach, Switzerland), DNA amplification and processing was performed with reagents from Promega (Dübendorf, Switzerland).

### Trypanosome cultures

Bloodstream form TbCls KO parasites were cultured at 37 °C in HMI-9 containing 15% (v/v) heat-inactivated fetal bovine serum, 1 μg/ml G418, 0.5 μg/ml hygromycin, 0.1 μg/ml puromycin, 1 μg/ml blasticidin (InvivoGen, Muttenz, Schweiz), and in the presence or absence of 1 μg/ml tetracycline to maintain or ablate, respectively, TbCls. TbCls KO parasites expressing *in-situ*-tagged proteins were cultured in the presence of an additional 1.5 μg/ml phleomycin.

### Generation of TbCls conditional knock-out mutants

To generate plasmids to replace the endogenous TbCls genes, blasticidin and hygromycin resistance cassettes consisting of a PARP promoter, the resistance gene and a tubulin 3’ UTR were amplified from pXS2 expression vectors (courtesy of James D. Bangs, University of Buffalo, NY) and inserted into a plasmid containing 400 bp long recombination sequences flanking the TbCls ORF (Serricchio & Bütikofer, 2012). Prior to transfections into NY single marker bloodstream forms (Wirtz, Leal, Ochatt, & Cross, 1999), plasmids were cut upstream and downstream of the recombination sequences using *XhoI* and *NotI*. The inducible hemagglutinin (HA)-tagged ectopic copy of TbCls was constructed by PCR amplification of the Tb927.4.2560 ORF as described (Serricchio & Bütikofer, 2012). Prior to transfection, the vector was linearized with *Not*I. Clones were obtained by limiting dilution and antibiotic selection using 1 μg/ml blasticidin, 0.1 μg/ml puromycin, 0.5 μg/ml hygromycin and 1 μg/ml tetracycline. Clones were PCR-tested for correct integration using primer 5’UTR_control TCGTCCGCGCCTTTGTGTAGCTA, which binds 50 bp upstream of the 5’-recombination site, in combination with different reverse primers. Construction of c-Myc-tagged proteins was done as described (Schädeli, et al., 2019).

### Southern blot analysis

For Southern blot analysis, 1.5 μg *Nco*I or *Pst*I/*Sph*I-digested genomic DNA was separated on a 1% agarose gel and transferred to hybond-N+ nylon transfer membrane (Amersham Pharmacia Biotech, Glattbrugg, Switzerland) using 10xSSC buffer (150 mM Na3-citrate, pH 7.0, 1.5 M NaCl) as described (Serricchio & Bütikofer, 2012). The membrane was probed with a 400 bp ^32^P-labelled PCR product of the TbCls 3’UTR generated with the prime-a-gene labelling system (Promega). Primers used for PCR were: 3’UTR_fwd CCCTCTAGACAGCTCACGAACCGTGCCCTA and 3’UTR_rev: CCCGCGGCCGCTATCCGTCGAGGGCCACCC. Hybridized probe was detected by autoradiography using BioMax MS films in combination with intensifying screens.

### Metabolic labeling

Approximately 10^7^ parasites in mid-log phase were labelled with 10 μCi [^3^H]-glycerol for 6 h followed by washing, lipid extraction, thin-layer chromatography and radioisotope scanning as described (Serricchio & Bütikofer, 2013).

### Preparation of crude membrane fractions and membrane proteins

Crude mitochondrial preparations were obtained by digitonin extraction as described elsewhere (Charriere, Helgadottir, Horn, Soll, & Schneider, 2006). Briefly, 10^8^ trypanosomes were washed in SBG buffer (150 mM Tris·HCl, pH 7.9, 20 mM glucose monohydrate, 20 mM NaH_2_PO_4_) and parasites collected by centrifugation (1500 x g, 5 min). Cells were suspended in 0.5 ml SoTE (20 mM Tris·HCl, pH 7.5, 0.6 M sorbitol, 0.2 mM EDTA) followed by the addition of SoTE containing 0.05% (w/v) digitonin. After 5 min incubation on ice, non-lysed cells were removed by centrifugation for 5 min at 800 x g and crude membrane fraction was collected by another centrifugation step (6000 x g, 5 min, 4° C). For complete membrane lysis and extraction of membrane proteins, the pellet was dissolved in 100 μl extraction buffer (20 mM Tris·HCl, pH 7.2, 15 mM NaH_2_PO_4_, 0.6 M sorbitol) containing 1.5% (w/v) digitonin and incubated on ice for 15 min. Solubilized proteins were cleared from insoluble material by centrifugation (16’000 x g, 30 min at 4 °C) and used for further analysis.

### Immunoprecipitation of TbCls-HA

To immunoprecipitate TbCls-HA, crude membranes isolated from 5×10^8^ parasites were solubilized with 100 μl RIPA buffer (25 mM Tris-HCl, pH 7.4, 150 mM NaCl, 0.1% SDS, 0.5% sodium deoxycholate, 1% NP-40) and heated to 65 °C for 5 minutes. After dilution with 900 μl IP buffer (10 mM Tris-HCl, pH 7.4, 150 mM NaCl, 1 mM EDTA, 1% Triton X-100, 0.5% NP-40, protease inhibitors) and centrifugation (16’000 x g, 30 min at 4 °C), anti-HA (16B12, Enzo Life Sciences) antibody was added in combination with Protein G Dynabeads (Thermo Fisher Scientific) and incubated for 16 h. After washing with IP buffer, proteins were eluted by SDS sample buffer and analyzed by immunoblotting as described below.

### Polyacrylamide gel electrophoresis and immunoblotting

Proteins were denatured by SDS and separated using 12% polyacrylamide gels under reducing conditions (SDS-PAGE). Alternatively, blue native polyacrylamide gel electrophoresis (BN-PAGE) was performed using digitonin extracts and separation by 4-12% acrylamide gradient gels at 4 °C (Wittig, Braun, & Schagger, 2006). Subsequently, proteins were transferred onto nitrocellulose membranes (Thermo Scientific) or Immobilon-P polyvinylidene difluoride membranes (Millipore, Billerica, MA) using a semi-dry protein blotting system (BioRad, Cressier, Switzerland). After blocking the membrane in TBS (10 mM Tris·HCl pH 7.5, 144 mM NaCl) containing 5% (w/v) milk powder, membranes were exposed to primary antibodies mouse anti-Hsp70 (provided by André Schneider, University of Bern, Switzerland or (Panigrahi et al., 2008)), mouse anti-HA (HA.11, 16B12, Enzo Life Sciences, Lausen, Switzerland), rabbit anti-ATP synthase subunits β, p18, ATPaseTb1, ATPaseTb2 and OSCP (Subrtova, et al., 2015), rabbit anti-AAC and anti-VDAC (Singha, Sharma, & Chaudhuri, 2009), mouse anti-AOX (provided by Minu Chaudhuri, Chicago Medical School, Chicago, IL), diluted 1:1000–1:5000 in TBS containing 5% (w/v) milk powder. Horseradish peroxidase-conjugated secondary anti-mouse and anti-rabbit antibodies (Dako, Glostrup, Denmark) were used at concentrations of 1:5000 and 1:1000, respectively, and detected using an enhanced chemiluminescence detection kit (Thermo Scientific). Protein sizes were determined using PageRuler Plus Prestained Protein Ladder (Thermo Scientific) and NativeMark™ Unstained Protein Standard (Invitrogen). For protein quantitation, total cell lysates from 4×10^6^ cells were loaded on TGX stain-free precast gels (BioRad) and subjected to SDS-PAGE before transfer to polyvinylidene difluoride membranes. Subsequently, proteins were immunodetected by specific antibodies and visualized using ChemiDoc™ Gel Imaging System. Signals from TbCls-depleted cells were compared to control samples and then normalized to Hsp70 loading control. The relative expression of the individual proteins was plotted using Graph Pad Prism 8.2.1.

### Transmission electron microscopy

TbCls KO parasites were cultured in the presence or absence of tetracycline to maintain or induce, respectively, ablation of TbCls expression. Trypanosomes were washed in PBS (pH 7.4, 137 mM NaCl, 2.7 mM KCl, 10 mM Na_2_HPO_4_, 1.8 mM KH_2_PO_4_) and processed for TEM analysis as described elsewhere (Dawoody Nejad, Serricchio, Jelk, Hemphill, & Bütikofer, 2018; Schädeli, et al., 2019).

### Stable isotope labeling with amino acids in cell culture (SILAC)

SILAC and liquid chromatography-mass spectrometry/mass spectrometry was done as described before (Schädeli, et al., 2019).

### MitoTracker staining

Live trypanosomes (2×10^6^ cells) were stained in culture medium with 100 nM MitoTracker Red CM-H_2_XRos (Invitrogen) for 30 min. After washing, parasites were resuspended in PBS, allowed to adhere to a microscope slide (Thermo Scientific) for 15 min, fixed in PBS containing 4% (w/v) paraformaldehyde for 10 min, washed and air-dried before mounting with Vectashield (Vector Laboratories, Burlingame, CA) containing 1.5 μg/ml 4’,6-diamidino-2-phenylindole (DAPI). The images were acquired using a Leica DMI6000 B microscope with 60x oil objective.

### Metabolomic analysis

Metabolites were extracted from 10^8^ parasites by rapid cooling to 4 °C by submersion of the tube in a dry ice/ethanol bath. After centrifugation for 10 minutes at 1000 x g, the supernatant was removed completely and the pellet suspended in 200 μl chloroform/methanol/water (1:3:1 ratio) at 4 °C. After mixing with a pipette, samples were rocked for 1 h at 4 °C, cleared at 13000 x g for 3 min and 180 μl of the supernatant was transferred into a new tube and stored at −80 °C until analysis.

Hydrophilic interaction liquid chromatography (HILIC) was carried out on a Dionex UltiMate 3000 RSLC system (Thermo Fisher Scientific, Hemel Hempstead, UK) using a ZIC-pHILIC column (150 mm × 4.6 mm, 5 μm column, Merck Sequant). The column was maintained at 30 °C and samples were eluted with a linear gradient of solvent A (20 mM ammonium carbonate in water) in acetonitrile over 26 min at a flow rate of 0.3 ml/min. For mass spectrometry (MS) analyses, a Thermo Orbitrap Fusion (Thermo Fisher Scientific) was operated in polarity switching mode and the MS settings were as follows: resolution 120’000; AGC 2e5; m/z range 70–1000; sheath gas 40; Auxiliary gas 5; sweep gas 1; probe temperature 150 °C; capillary temperature 325 °C. For positive mode ionization: source voltage +4.3 kV. For negative mode ionization: source voltage −3.2 kV. S-Lens RF level 30%. Fragmentation was performed with the following parameters: collision energy: 60%; stepped collision energy: 35%; isolation window: 2; dynamic exclusion after 1 time; exclusion duration: 6 seconds; exclude isotopes: true; minimum intensity: 50,000. Instrument raw files were converted to positive and negative ionization mode mzXML files. These files were then analyzed using PiMP (Gloaguen, et al., 2017) in combination with FrAnK (an in-house fragmentation tool).

### Mitochondrial membrane potential (Δψm)

*In vivo* Δψm was measured using the cell-permeant red-fluorescent dye TMRE (tetramethylrhodamine ethyl ester, Thermo Fisher Scientific). For each time point, an equal number of parasites (3×10^6^) was harvested and resuspended in culture medium containing 60 nM TMRE. Mitochondrial staining was carried out for 30 min under standard culture conditions (37 °C and 5% CO_2_). Subsequently, trypanosomes were spun down at 1400 x g for 10 min at room temperature, resuspended in 1 ml of 1x PBS (10 mM phosphate buffer, 130 mM NaCl, pH 7.3), and immediately analyzed by flow cytometry (BD FACS Canto II Instrument) using the PE filter. For each sample, 10’000 events were collected. Treatment with 20 μM FCCP (carbonyl cyanide 4-(trifluoromethoxy) phenylhydrazone) was used as a control for mitochondrial membrane depolarization. Data were evaluated using BD FACS Diva (BD Company) software. The experiments were performed in triplicates.

*In situ* Δψm of permeabilized cells was determined fluorometrically employing safranin O. For each time point, 2×10^7^ parasites were harvested and washed once with ANT buffer (8 mM KCl, 110 mM K-gluconate, 10 mM NaCl, 10 mM free-acid Hepes, 10 mM K_2_HPO_4_, 0.015 mM EGTA potassium salt, 10 mM mannitol, 0.5 mg/ml fatty acid-free bovine serum albumin, 1.5 mM MgCl_2_, pH 7.25) (Chinopoulos et al., 2009). The cell pellet was resuspended in 200 μl of ANT buffer containing 5 μM safranin O, 40 μM digitonin and 2 mM ATP (PanReac AppliChem), and subsequently transferred into a white flat-bottom 96-well microtiter plate. Fluorescence was recorded in a Tecan Infinite^®^ 200 PRO series plate reader using 496 and 586 nm excitation and emission wavelengths, respectively. The F_o_F_1_-ATP synthase inhibitor oligomycin (10 μg/ml) and the uncoupler SF6847 (250 nM) (Enzo Life Sciences) were added where indicated. The experiments were performed in triplicates.

### Oxygen flux analysis

The oxygen consumption rate was determined using the Oroboros Oxygraph-2K (Oroboros Instruments Corp., Innsbruck, Austria). For each time point, 2×10^7^ parasites were harvested and washed once with Mir05 mitochondrial respiration medium (0.5 mM EGTA, 3 mM MgCl_2_, 60 mM lactobionic acid, 20 mM taurine, 10 mM KH_2_PO_4_, 20 mM Hepes, 110 mM sucrose, 1 mg/ml fatty acid-free bovine serum albumin, pH 7.1). The pellet was resuspended in 2.1 ml of Mir05 and transferred into the respiration chamber at 37 °C under constant stirring. To trigger AOX-mediated respiration, 20 mM DL-glycerol-3-phosphate was added. Once the maximal respiration rate was achieved, respiration was inhibited by addition of 250 μM SHAM (salicylhydroxamic acid). The most stable portion of either the oxygen consumption rate slope or the oxygen concentration in the chamber slope was determined for each biological replicate after the addition of the substrate and the inhibitor. The values were plotted and analyzed statistically using GraphPad Prism 8.0 software.

### ATP measurements

Both the ADP/ATP ratio and the total cellular ATP content were measured using a bioluminescence-based ADP/ATP assay kit (Sigma) following the manufacturer’s protocol. In brief, 1×10^6^ parasites per time point were harvested and washed once with PBS-G (1x PBS containing 6 mM glucose). Cells were resuspended in 10 μl of PBS-G and transferred into a white flat-bottom 96-well microtiter plate. Bioluminescence was recorded using an Orion II microplate luminometer (Titertek Berthold) and the ADP/ATP ratios were calculated according to the manufacturer’s protocol. The ATP content (first fluorescence read of the assay) of TbCls-depleted trypanosomes was expressed relative to control parasites using GraphPad Prism 8.0 software.

## Acknowledgements

The work was supported by grant no. 169355 from the Swiss National Science Foundation to P.B, and by Czech Science Foundation (18-17529S) and ERD fund (CZ.02.1.01/0.0/0.0/16_019/0000759) to A.Z. We thank André Schneider (University of Bern) and Minu Chaudhuri (Chicago Medical School) for antibodies and Anant K. Menon (Weill Cornell Medical College New York) for advice. P.B. thanks A. Niederer, M. Bütikofer and R. Plant for support.

## Conflict of interest

The authors declare that they have no conflicts of interest with the contents of this article.

## Author contributions

MS, DS, AZ and PB designed research; MS, CHY, DS, HBH, AH and JG performed research; MS, CHY, DS, AH, JG, AZ and PB analyzed data; MS, AZ and PB wrote the manuscript. All authors reviewed the manuscript.

## Notes

### Competing Interest Statement

The authors have declared no competing interest.

